# T cell transcriptional landscapes are shaped by TCR sequence similarity

**DOI:** 10.1101/2023.11.27.568912

**Authors:** Hao Wang, Zhicheng Ji

## Abstract

T cells exhibit high heterogeneity in both their gene expression profiles and antigen specificities, but the precise relationship between these two features remains largely unknown. Utilizing a large-scale single-cell immune-profiling dataset that includes both T cell receptor (TCR) sequences and gene expression profiles of individual cells, we systematically investigated how the transcriptional profiles of T cells are linked to their TCR sequence similarity. We observed that T cells sharing the same clonotype often belong to the same cell type, and that gene expression similarity among T cells correlates with TCR sequence similarity. Furthermore, we identified features such as clonality, VJ gene usage, sequence motifs, and amino acid preferences that are distinctly associated with specific T cell subtypes. These findings suggest that T cell transcriptional landscapes are broadly shared and influenced by TCR sequence similarity.

## Introductions

T cells are a fundamental component of adaptive immunity. The function of T cells is largely determined by the specificity of their T cell receptors (TCRs) and their gene expression programs. Understanding the association between TCR sequences and gene expression is crucial for elucidating how antigen specificity is linked to T cell activation, differentiation, and effector functions. Prior studies have found that T cells from the same clonotype often exhibit similar gene expression profiles in various contexts, including lung, liver, and colorectal cancers^1–4^, breast tumors^5^, yellow fever^6^, and homeostasis^7^. These findings suggest that TCR specificity may shape transcriptional programs in both health and disease. However, the precise relationship between TCR sequence diversity and transcriptional heterogeneity remains poorly understood.

The advent of single-cell immune profiling has enabled researchers to explore the relationship between TCR sequences and gene expression with unprecedented resolution. This approach allows for the direct integration of TCR sequence diversity with transcriptional states, providing valuable insights into how different clonotypes correspond to the functional states of T cells. Several computational methods, such as CoNGA^8^, tessa^9^, and mvTCR^10^, have been developed to integrate TCR sequence information with gene expression, offering a systematic approach to studying this association. Understanding how TCRs are linked to gene expression not only deepens our knowledge of T cell-mediated immunity but also facilitates the development of more effective computational tools for analyzing single-cell immune profiling data.

In this study, we leveraged the human Antigen Receptor database (huARdb)^11^, a large single-cell immune profiling database, to systematically explore the association between gene expression and TCR sequence similarities. The database includes a total of 2,228,532 *αβ* T cells collected from 972 samples and 559 individuals. For each T cell, both gene expression and TCR sequences were measured simultaneously. While this database is well-suited for studying the association between TCRs and the transcriptional profiles of T cells, a systematic analysis remains lacking. We found that T cells with similar CDR3 sequences also exhibit similar gene expression profiles across different samples and cell types. Additionally, we identified specific CDR3 motif sequences and VJ gene usages that are enriched in particular cell types. Our findings reveal that the transcriptional profiles of T cells are largely associated with and shaped by T cell clonotypes, suggesting a functional connection between antigen specificity and cellular state.

## Results

### T cells of the same clonotype exhibit similar transcriptional profiles

We begin by investigating the gene expression similarity of T cells that belong to the same clonotype and originate from the same sample. In this study, a clonotype is defined as a group of T cells that share identical amino acid sequences in both their alpha and beta chains. Only clonotypes containing multiple T cells were analyzed. Gene expression similarity is examined at three nested levels (Methods). The first level assesses whether all T cells within a clonotype are CD4+ or CD8+ by calculating the purity of the most abundant T cell type (Figure 1a). The second level examines whether CD4+ or CD8+ T cells within a clonotype belong to the same subtype (e.g., CD4+ Treg) by calculating the purity of the most abundant T cell subtype (Figure 1a). The third level evaluates gene expression similarity among T cells of the same subtype by measuring their distances in a PCA space based on gene expression profiles (Figure 1b).

**Figure 1.**
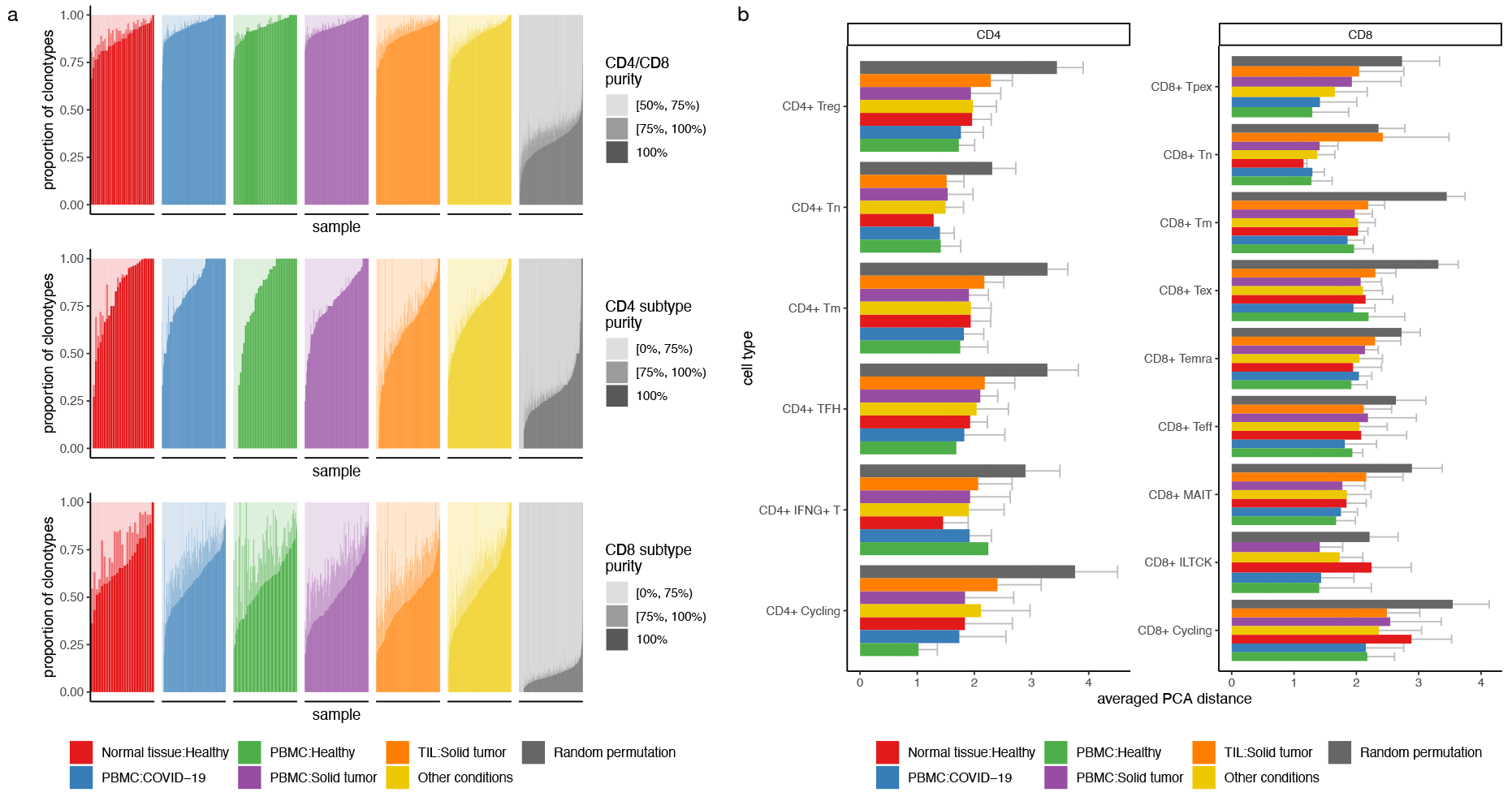
Gene expression similarity of T cells from the same clonotype. **a**, Proportion of clonotypes with varying levels of CD4/CD8 purity, CD4 subtype purity, and CD8 subtype purity, from top to bottom. Each column represents one sample. Samples from different conditions are colored differently. For the random permutation, samples from all conditions were pooled together. **b**, PCA distances for clonotypes within a sample were averaged and shown as individual data points. Each bar represents the distribution of averaged PCA distances for samples from one condition within one T cell subtype. Samples from different conditions are colored differently. For the random permutation, samples from all conditions were pooled together. CD4 and CD8 T cell subtypes are displayed in two separate panels. The error bars represent standard deviations.

Across all samples, an average of 90.0% of clonotypes consist of T cells that all belong to either CD4+ or CD8+ T cells. Additionally, an average of 92.9% of clonotypes have more than 75% of their cells belonging to the same T cell type. Among CD4+ T cells, an average of 74.1% of clonotypes consist of T cells that belong to the same CD4+ T cell subtype, while 75.9% of clonotypes have more than 75% of their cells belonging to the same CD4+ T cell subtype. Similarly, among CD8+ T cells, an average of 55.6% of clonotypes consist of T cells that belong to the same CD8+ T cell subtype, and 66.8% of clonotypes have more than 75% of their cells belonging to the same CD8+ T cell subtype. This pattern is consistent across samples collected from different tissues and from donors with varying disease statuses. Moreover, the observed purity is substantially higher than that obtained after randomly permuting cell type labels. Figure 1b further illustrates that PCA distances among cells of the same type are substantially lower than those observed after randomly permuting cell positions in PCA space.

Collectively, these results suggest that cells within the same clonotype exhibit highly similar gene expression profiles. This preserved transcriptional identity during clonal expansion is likely due to the shared progenitor origins and differentiation pathways of T cells within the same clonotype. Consistent patterns across tissues and disease statuses suggest that these mechanisms are robust.

### T cell clonotypes shared across individuals display unique transcriptional profiles

We then examined T cells that belong to the same clonotype but originate from different individuals. Such shared clonotypes have been reported in previous studies^12–14^, but a systematic analysis of their transcriptional profiles is lacking. There are 3,729 clonotypes whose T cells are found in multiple individuals (Figure 2a). The gene expression profiles of these shared clonotypes differ substantially from those of private clonotypes, whose T cells are found in a single individual (Figure 2b). For example, most T cells from shared clonotypes are CD8+ T cells. Within the CD8+ T cell population, T cells from shared clonotypes are more enriched in CD8+ Teff and less abundant in CD8+ Tm, CD8+ Tex, and CD8+ Tn. This suggests that shared clonotypes likely arise from immune responses to common pathogens, such as widespread viral infections, which drive transient effector expansion rather than long-term memory formation. In contrast, private clonotypes may reflect individual-specific antigen exposure, chronic stimulation, or stochastic TCR recombination.

**Figure 2.**
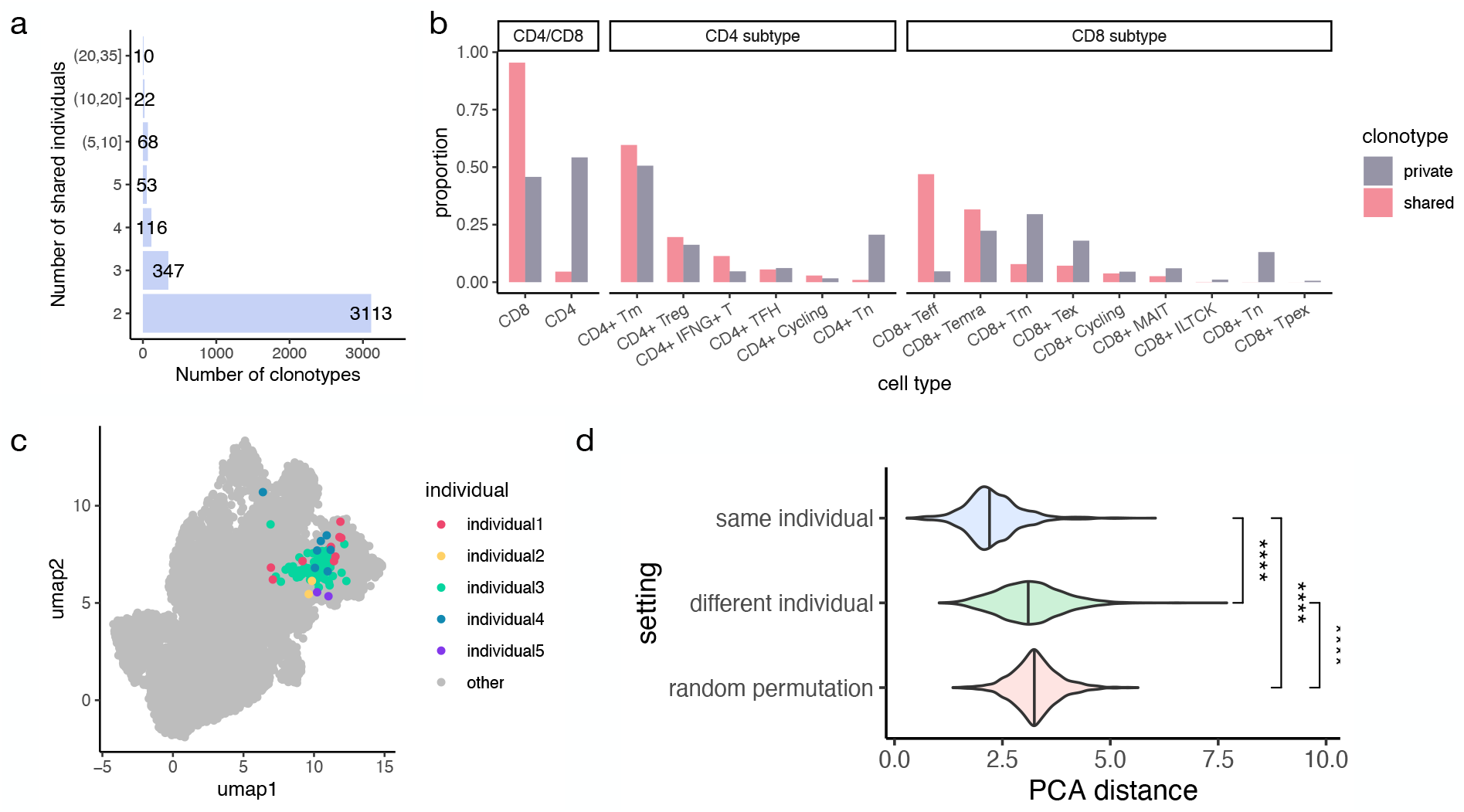
Gene expression profiles of T cell clonotypes shared across individuals. **a**, Number of clonotypes (x-axis) with T cells shared across different numbers of individuals (y-axis). **b**, Proportions of CD4 and CD8 T cells among all T cells, proportions of CD4 T cell subtypes among CD4 T cells, and proportions of CD8 T cell subtypes among CD8 T cells for private and shared clonotypes, from left to right. Private clonotypes contain T cells from only one individual, while shared clonotypes include T cells from multiple individuals. **c**, UMAP visualization of T cells from an example shared clonotype. Colors represent T cells from different individuals belonging to the clonotype. Grey indicates other T cells not belonging to the clonotype. **d**, Distribution of PCA distances for pairs of T cells: within the same individual and shared clonotype, between different individuals but the same shared clonotype, and after random permutation of cell coordinates in PCA space.

T cells from the same shared clonotype also show similarity in gene expression. Figure 2c visualizes the distribution of T cells from one example shared clonotype on the UMAP. Within each shared clonotype, we also calculated PCA distances between T cells from the same and different individuals (Figure 2d), which were further compared to the PCA distances when PCA coordinates were randomly permuted across T cells from all shared clonotypes. Consistent with the findings in the previous section, T cells from the same individual are close to each other. While T cells from different individuals are more distinct from those of the same individual, they are slightly more similar compared to the random permutation. These results suggest that T cells from the same clonotype preserve gene expression similarity to some extent, even when they originate from different individuals.

### T cell transcriptional similarity correlates with TCR similarity

We next studied the gene expression similarity among T cells from different clonotypes within each sample. Similar to the previous section, gene expression similarity was examined at three nested levels (Methods). The purity of CD4+ and CD8+ T cells shows a decreasing trend with increasing Damerau–Levenshtein distances in CDR3a or CDR3b sequences (Figure 3a). However, this decrease is only significant when the CDR3a/b sequence distance is less than 4, and the purity remains nearly unchanged when the distance exceeds 4. Within CD4+ and CD8+ T cells, the purity of CD4+ and CD8+ subtypes follows a similar decreasing trend (Figure 3a). Additionally, within each CD4+ and CD8+ cell subtype, PCA distance increases with increasing CDR3a/b sequence distances (Figure 3b). However, the cutoff point at which PCA distance stops increasing varies across cell subtypes. For example, CD4+ Tregs exhibit a nearly constant PCA distance when the CDR3a sequence distance is greater than 0, whereas CD8+ MAIT cells display a linearly increasing pattern when the CDR3a sequence distance is less than 7.

**Figure 3.**
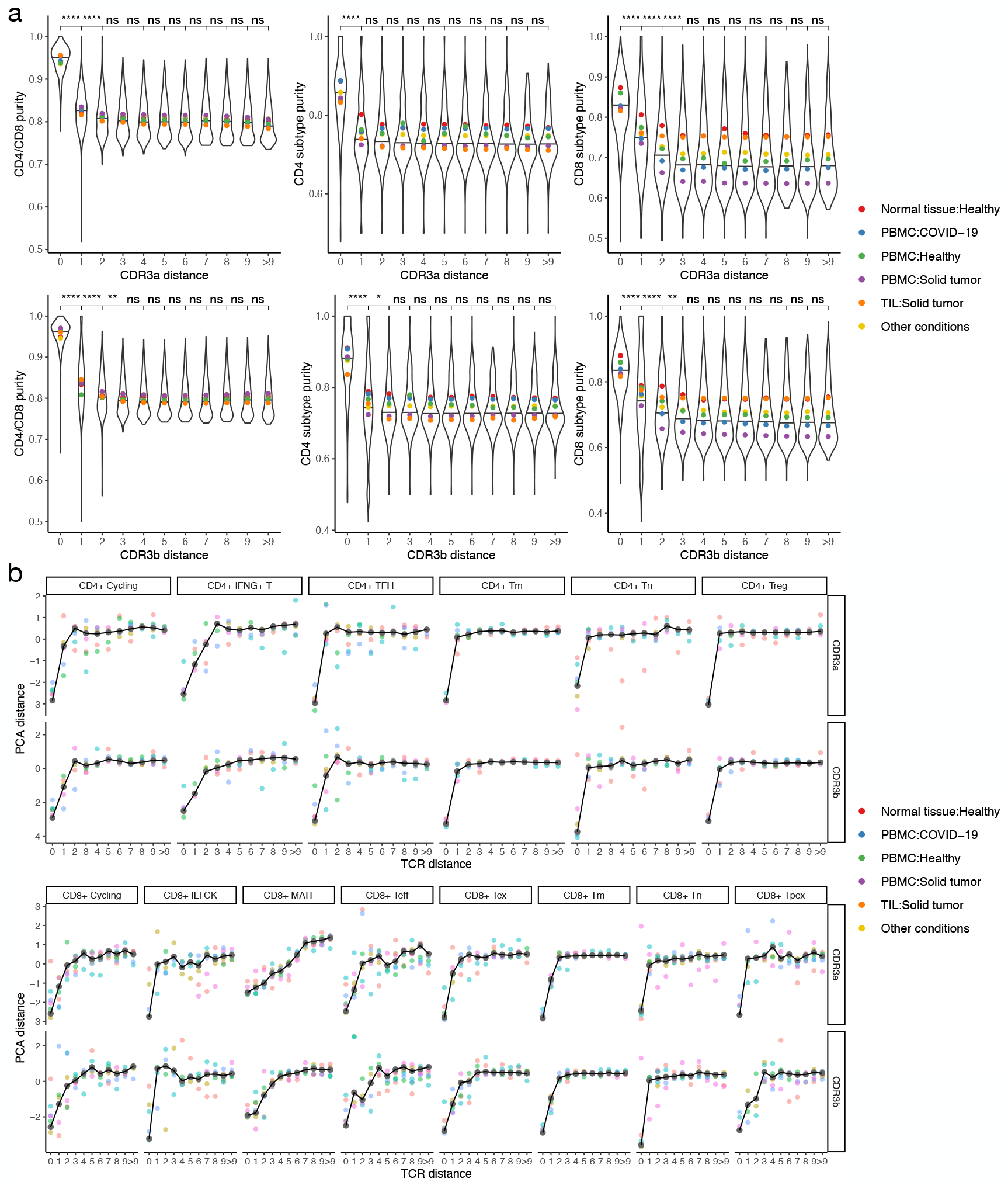
Association between gene expression similarity and CDR3 sequence similarity. **a**, Distribution of CD4/CD8 purity, CD4 subtype purity, and CD8 subtype purity (x-axis, from left to right) across different CDR3a and CDR3b distances (y-axis, from top to bottom). Each data point in the violin plot represents the average purity within a sample. Colored dots indicate the average sample-level purity across samples under specific conditions. **b**, Average PCA distance (y-axis) across different CDR3 distances (x-axis). Each black dot represents the PCA distance averaged across clonotype pairs from all samples. Lines connect the black dots. Colored dots represent the average PCA distance across clonotype pairs from samples under specific conditions.

These findings indicate that T cell gene expression is closely linked to clonotype similarity up to a specific threshold. When T cells have minor differences in their CDR3 sequences, their gene expression remains relatively similar. However, beyond a certain sequence divergence threshold, additional differences in clonotype do not appear to further influence gene expression. This transcriptional similarity could be attributed to the coordinated activities of T cells with similar antigen specificity, as the threshold aligns with the sequence similarity observed among clonotypes that recognize the same type of antigen^15^.

### T cell subtypes exhibit diverse clonality and sequence patterns

We then explored the similarity of TCR sequences within each T cell subtype by calculating clonality, which measures how evenly clonotypes are distributed (Methods). We found that CD8+ T cells exhibit higher clonality than CD4+ T cells (Figure 4a). While different CD4+ T cell subtypes have comparable clonality levels (Figure 4b), clonality varies substantially among CD8+ subtypes (Figure 4c). For example, CD8+ Teff and CD8+ Temra have substantially higher clonality than CD8+ MAIT and CD8+ Tm. This aligns with their functions, as CD8+ T cells undergo strong clonal expansion for targeted cytotoxic responses, whereas CD4+ T cells maintain a more diverse repertoire for immune regulation and coordination.

**Figure 4.**
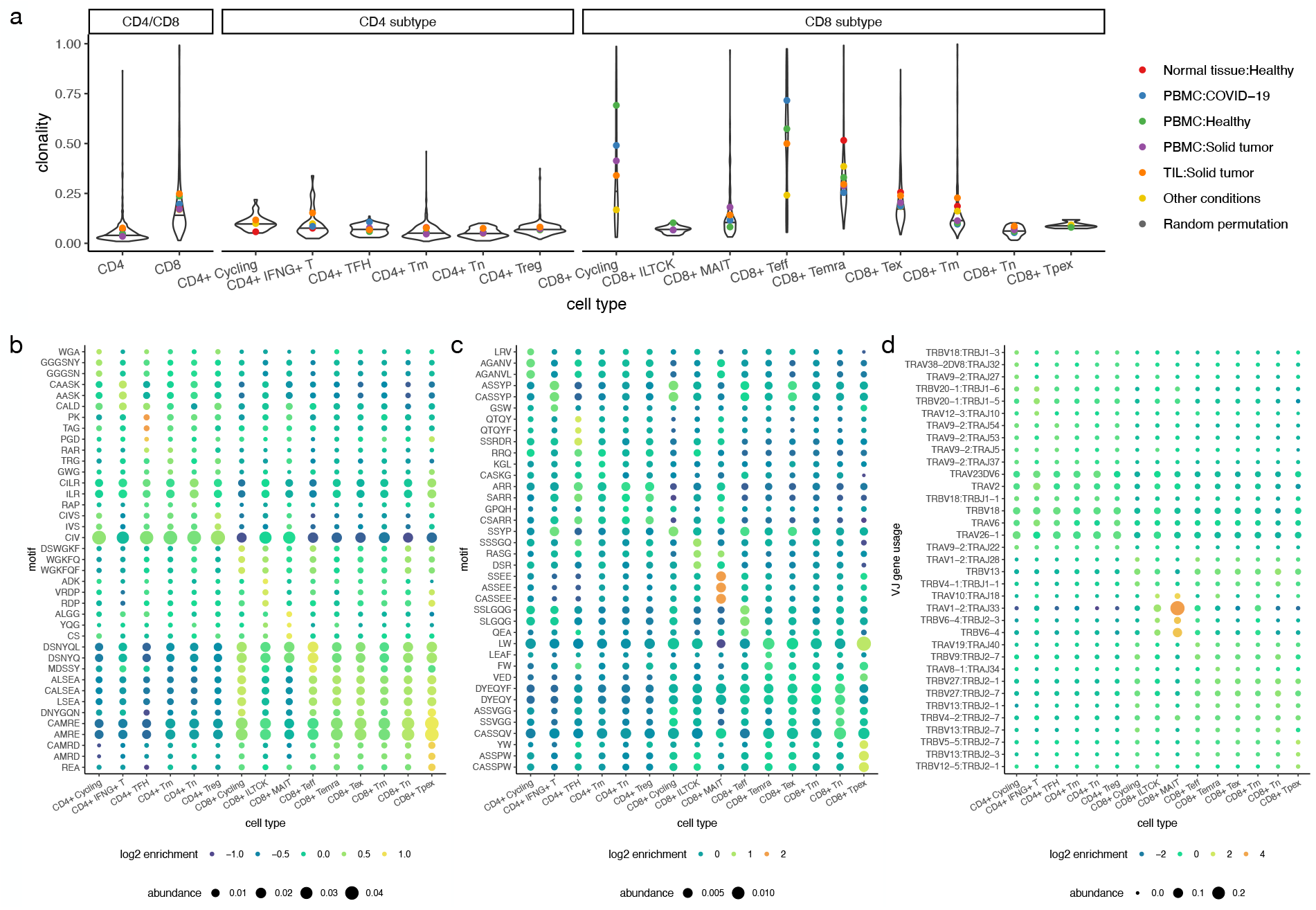
Analysis of clonality, motifs, and VJ gene usage. **a**, Distribution of clonality in CD4 and CD8 cells, CD4 T cell subtypes, and CD8 T cell subtypes, from left to right. Each data point in the violin plot represents the clonality value of one sample. Colored dots indicate the average sample-level clonality for samples under specific conditions. **b–c**, Enriched motifs (y-axis) in different T cell subtypes (x-axis) for CDR3a (**b**) and CDR3b (**c**). For each T cell subtype, the top 3 motifs with the highest enrichment scores were selected. The color of each dot represents the log_2_ enrichment score, and the size of each dot represents the proportion of cells in the T cell subtype whose CDR3a/b sequences contain the motif. **d**, Enriched VJ gene usage (y-axis) in different T cell subtypes (x-axis). For each T cell subtype, the top 3 VJ gene usages with the highest enrichment scores were selected. The color of each dot represents the log_2_ enrichment score, and the size of each dot represents the proportion of cells in the T cell subtype that use the corresponding VJ genes.

Previous studies have reported that TCRs sharing antigen specificity exhibit conserved motifs or global similarity in CDR3 sequences^15^. We conducted a similar analysis and identified VJ gene usages and short motif sequences enriched in each T cell subtype (Figure 4b-d, Methods). The analysis was performed on unique CDR3a/b sequences to avoid bias introduced by varying clonotype expansion levels. Among these, VJ gene usages and motifs associated with CD8+ MAIT cells showed the highest enrichment. This is expected, as MAIT cells often bear the TRAV1-TRAJ33 chain and harbor evolutionarily conserved TCRs^16^. Interestingly, we also identified motifs enriched in cell types other than CD8+ MAIT cells. For example, the motif “DSNYQ” in the CDR3a sequence is enriched in CD8+ Teff cells as well as other CD8+ subtypes. However, no VJ gene usage was highly enriched in non-MAIT cell types. These results suggest that CD8+ MAIT cells are likely the only T cell subtype with evolutionarily conserved TCRs, while other cell types may be shaped more by antigen-driven selection.

The enrichment of short motifs in non-MAIT cell types suggests that certain CDR3 sequence patterns may contribute to antigen recognition, even in the absence of specific VJ gene constraints. This could indicate a degree of convergent recombination, where different TCRs independently acquire similar functional features. Additionally, the presence of shared motifs across multiple CD8+ subtypes raises the possibility that some TCRs recognize common antigenic determinants.

### T cell subtypes exhibit distinct amino acid preferences

Finally, we examined whether certain types of amino acids are more enriched in specific T cell subtypes. We grouped cells based on the length of their CDR3a/b sequences. For each position in the CDR3a/b sequences, we calculated the proportion of occurrences for each T cell subtype and amino acid pair and compared it with the expected proportion using a contingency table (Methods). Figure 5 presents the top 20 most enriched and depleted T cell subtype–amino acid pairs in CDR3a and CDR3b sequences.

**Figure 5.**
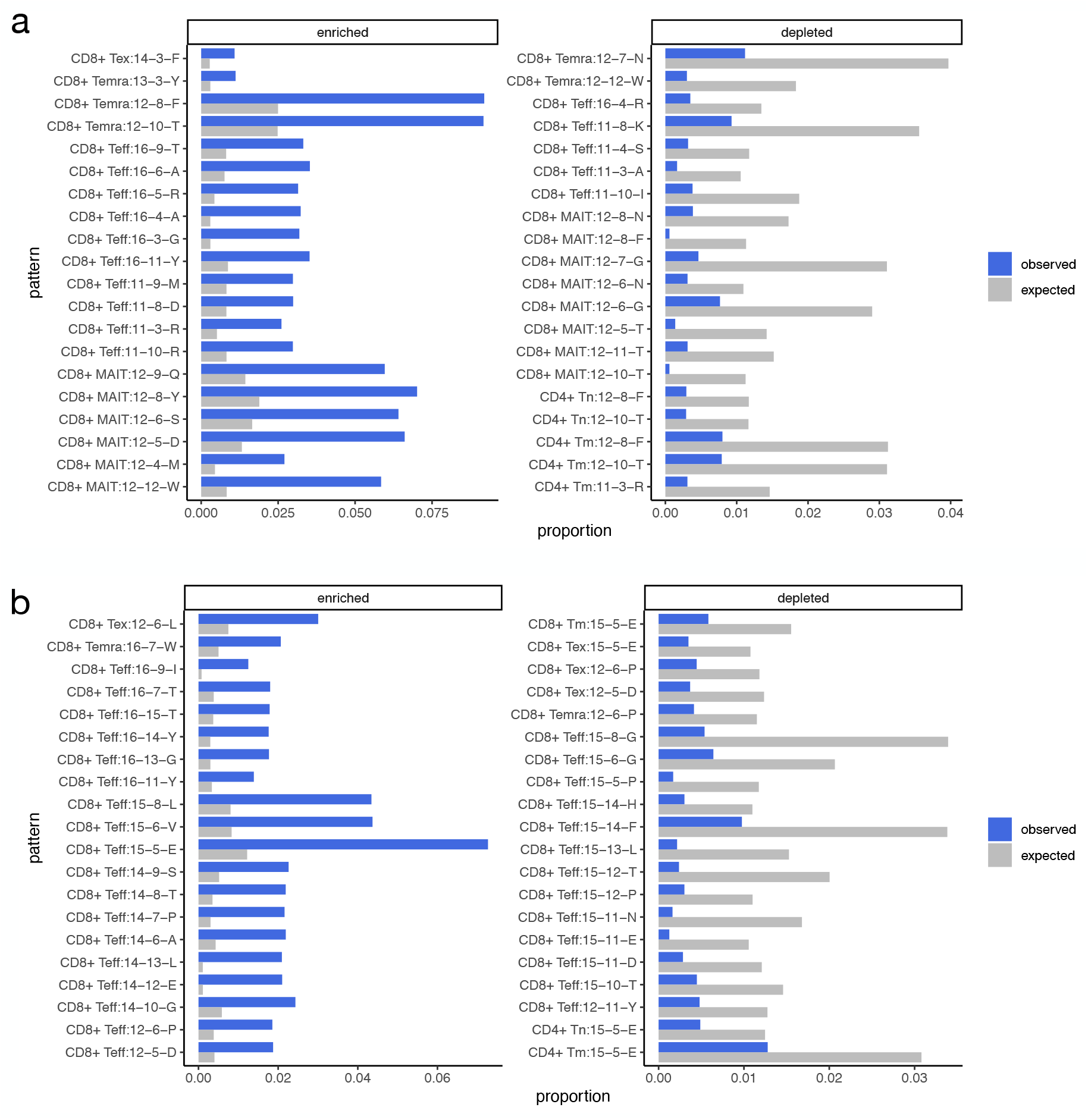
Amino acid preferences of T cell subtypes in CDR3a (**a**) and CDR3b (**b**). Each row represents a pattern, which is a combination of a T cell subtype and an amino acid at a specific position in CDR3a/b sequences of a given length. Each pattern is named using the T cell subtype, the full length of the CDR3a/b sequence, the position within the sequence, and the amino acid. For example, ‘CD8+ Tex:14-3-F’ in **a** refers to the CD8+ Tex subtype with amino acid F at the third position of CDR3a sequences of length 14. Observed and expected proportions, computed from contingency tables, are shown on the x-axis. Patterns with either observed or expected proportions greater than 1% were retained. The top 20 patterns with the highest O/E ratios were selected as the most enriched patterns for visualization. Likewise, the top 20 patterns with the lowest O/E ratios were selected as the most depleted patterns.

We found that the observed proportions of certain pairs were substantially higher than expected. For example, the pair CD8+ MAIT and amino acid tryptophan(W) at position 12 of the length-12 CDR3a sequence had an observed proportion of 5.84%, which is 7.05 times the expected proportion of 0.83%. This finding aligns with previous studies and reflects the evolutionarily conserved nature of MAIT TCRs. However, we also identified other enriched pairs associated with non-MAIT cell types. For instance, the pair CD8+ Teff and amino acid alanine(A) at position 4 of the length-16 CDR3a sequence had an observed proportion of 3.23%, which is 10.91 times the expected proportion of 0.30%. This highlights potential functional constraints on TCR recognition and may provide insights into how different T cell subsets recognize antigens and mediate immune responses.

## Conclusions

In this study, we systematically explored the relationship between gene expression and TCR sequences using data from single-cell immune profiling. We found that T cell transcriptional landscapes are largely determined by the similarity of their TCR sequences. Specifically, we found that T cells with the same clonotype exhibit similar gene expression profiles and that T cell gene expression similarity correlates with TCR sequence similarity. Additionally, we identified clonality, VJ gene usage, motif sequences, and amino acid preferences that are distinctly associated with T cell subtypes.

Our findings suggest that T cell transcriptional states are not solely dictated by external stimuli or differentiation stages but are intrinsically linked to the specificity of their TCR sequences. The strong correlation between TCR sequence similarity and gene expression profiles implies that antigen recognition and clonal expansion may drive distinct transcriptional programs, reinforcing the idea that TCR signaling plays a direct role in shaping T cell fate. Moreover, our findings highlight the potential for leveraging TCR sequence features to predict functional states. This suggests that TCR-based classification could enhance our understanding of T cell heterogeneity and improve immune repertoire analysis in various contexts, including infection, autoimmunity, and cancer immunotherapy. Future studies should explore whether these transcriptional patterns are conserved across different immune conditions and how they evolve in response to antigen exposure over time.

## Methods

### Dataset

We downloaded information on TCR CDR3a/b amino acid sequences, cell types, sample metadata, processed gene expression matrix, and low dimensional representations of PCA and UMAP from huARdb version 2.0.0a (https://huarc.net/v2/). Details of the data processing procedure can be found on the huARdb website and in the original publication^11^. The dataset contains 2,228,532 *αβ* T cells, of which 51.3% are CD4+ T cells and 48.7% are CD8+ T cells. There are six CD4+ T cell subtypes and nine CD8+ T cell subtypes. Each cell in the database contains exactly one CDR3a and one CDR3b amino acid sequence. A clonotype is defined as a group of T cells with identical CDR3a and CDR3b amino acid sequences. The cells were collected from 972 samples and 559 individuals. Five conditions have more than 50 samples: Normal tissue (Healthy), PBMC (COVID-19), PBMC (Healthy), PBMC (Solid tumor), and TIL (Solid tumor).

### Analyzing T cells of a clonotype in a sample (Figure 1)

#### CD4/CD8 purity

For each clonotype with at least two cells in a sample, we computed the proportions of CD4+ and CD8+ T cells among all T cells in the clonotype. CD4/CD8 purity is defined as the higher of the two proportions. To obtain a background distribution of CD4/CD8 purity, we randomly permuted the labels of CD4 and CD8 T cells and recalculated CD4/CD8 purity using the same procedure. The analysis was performed only on samples with at least 500 cells and an overall CD4/CD8 purity of less than 0.9.

#### CD4/CD8 T cell subtype purity

For each clonotype with at least two CD4+ T cells in a sample, we computed the proportion of each CD4+ T cell subtype among all CD4+ T cells in the clonotype. CD4+ T cell subtype purity is defined as the highest of these proportions. Clonotypes that do not contain CD4+ T cells were ignored. To obtain a background distribution of CD4+ T cell subtype purity, we randomly permuted the labels of CD4+ T cell subtypes within CD4+ T cells and recalculated CD4+ T cell subtype purity using the same procedure. The analysis was performed only on samples with at least 500 CD4+ T cells and an overall CD4+ T cell subtype purity of less than 0.9. CD8+ T cell subtype purity was computed in a similar manner.

#### PCA distance

For each T cell subtype, we retained clonotypes that have at least two cells belonging to that T cell subtype. Within each clonotype, Euclidean distance between all possible pairs of cells that belong to the same T cell subtype were calculated on the PCA space that has 10 PCs. To obtain a background distribution of PCA distances, we randomly permuted the PCA coordinates of T cells within each T cell subtype and then recalculated the PCA distances using the same procedure. The analysis was performed only on samples with at least 500 cells.

### Analyzing T cells of a clonotype across multiple samples (Figure 2)

For each shared clonotype containing cells from multiple individuals, we computed the Euclidean distance between all possible pairs of cells within the clonotype. If a clonotype contains more than 1,000 cells, we randomly selected 1,000 cells for the computation. We also recorded whether each pair of cells belong to the same or different individuals. To obtain a background distribution of PCA distances, we randomly permuted the PCA coordinates across T cells pooled from all shared clonotypes and then recalculated the PCA distances using the same procedure.

### Analyzing T cells of different clonotypes in a sample (Figure 3)

#### CDR3 sequence distances

For each pair of unique CDR3a sequences, the sequence distance was calculated using optimal string alignment (the `stringdist` function in the R `stringdist` package). The sequence distance for each pair of unique CDR3b sequences was calculated similarly.

#### CD4/CD8 purity

CD4/CD8 purity was calculated for all pairs of unique CDR3a sequences within a sample. We designed a metric to account for differences in the number of elements between the two groups of cells that possess the two CDR3a sequences. Specifically, for a given pair of unique CDR3a sequences, let *v*_1_ be a vector of length *n*_1_, where each entry *v*_1,*i*_ is a CD4/CD8 label corresponding to a cell in the first group, indicating whether the cell belongs to the CD4+ or CD8+ T cell populations. Similarly, let *v*_2_ be a vector of length *n*_2_, where each entry *v*_2,*i*_ is a CD4/CD8 T cell label corresponding to a cell in the second group. To account for differences in group sizes, we repeated *v*_1_ *n*_2_ times and *v*_2_ *n*_1_ times. We then combined the two repeated vectors to form *v*. Finally, we computed the proportions of CD4 and CD8 labels among all labels in *v*, and CD4/CD8 purity was defined as the higher of the two proportions. To calculate CD4/CD8 purity when the CDR3a distance is 0, both *v*_1_ and *v*_2_ were set to the CD4/CD8 T cell labels for a group of T cells sharing the same CDR3a sequence, and the computation was performed as described above. The same procedure was applied to calculate CD4/CD8 purity for CDR3b sequences.

If there are more than 10,000 pairs of clonotypes with the same CDR3a sequence distance, we randomly sampled 10,000 pairs from all such pairs. Pairs of clonotypes with a CDR3a sequence distance exceeding 9 were grouped together before performing random sampling. The same random sampling procedure was performed when considering CDR3b sequence distances.

#### CD4/CD8 T cell subtype purity

To calculate CD4 T cell subtype purity, we focused only on CD4+ T cells. Purity was calculated in the same way as CD4/CD8 purity, except that CD4/CD8 T cell labels were replaced with CD4+ T cell subtype labels. Random sampling was performed within CD4 T cells, following the procedure described previously. CD8 T cell subtype purity was obtained using a similar procedure.

#### PCA distance

Within a sample, within a T cell subtype, and for each pair of unique CDR3a sequences, Euclidean distances were calculated between all pairs of cells where one cell possesses one CDR3a sequence and the other possesses the other. In cases where the CDR3a sequence distance was 0, Euclidean distances were calculated between pairs of cells sharing the same CDR3a sequence. Random sampling was performed within each T cell subtype, following the procedure described previously. The same computation was applied to CDR3b sequences.

### Calculation of clonality (Figure 4)

Simpson’s clonality was calculated within each sample and within CD4+ T cells, CD8+ T cells, or each T cell subtype. Specifically, denoting *p*_*i*_ as the proportion of cells belonging to clonotype *i*, Simpson’s clonality is defined as: 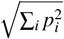.

### Motif and VJ gene usage analysis (Figure 4)

To perform motif analysis, each unique CDR3a sequence was split into consecutive 2-mers, 3-mers, 4-mers, 5-mers, and 6-mers with a stride of 1. Each k-mer was treated as a motif. For each CDR3a sequence, all motifs were pooled together, and duplicate elements were discarded. Motifs present in more than 1,000 cells were retained. Within each T cell subtype, the proportion of unique CDR3a sequences containing each motif was computed. To obtain a background distribution of motif proportions, we randomly permuted the T cell subtype labels across all cells and repeated the computation. For each motif, an enrichment score was calculated as (*p* + 0.001)*/*(*q* + 0.001) where *p* and *q* represent the proportions of the motif in the observed and permuted data, respectively. The same procedure was applied to CDR3b sequences.

VJ gene usage analysis was performed in a similar manner. Cells that share the same CDR3a or CDR3b sequence may exhibit different VJ gene usage. Therefore, for each unique clonotype, the sets of TRAV, TRAJ, TRBV, TRBJ, the TRAV–TRAJ pair, and the TRBV–TRBJ pair used by cells within the clonotype were identified, and duplicate elements were removed. The computation of proportions and enrichment scores was carried out in the same way as in the motif analysis.

### Analyzing amino acid preferences in T cell subtypes (Figure 5)

Cells with the same number of amino acids in their CDR3a sequences were grouped together. Groups with more than 100,000 cells were retained. Within each group, and for each amino acid position in the CDR3a sequence, a contingency table was constructed, where rows represent different amino acid choices at that position and columns represent the T cell subtypes of the cells. Only amino acids associated with more than 1,000 cells were included in the contingency table. Denote *n*_*i j*_ as the number of cells associated with the *i*th row and *j*th column in the contingency table. The observed and expected proportions of the element in the *i*th row and *j*th column of the contingency table were calculated as *n*_*i j*_*/* ∑_*a*_ ∑_*b*_ *n*_*ab*_ and (∑_*a*_ *n*_*aj*_)(∑_*b*_ *n*_*ib*_)*/*(∑_*a*_ ∑_*b*_ *n*_*ab*_)^2^, respectively. The observed to expected (O/E) ratio was calculated as the ratio between the observed and expected proportions for each element in the contingency table.

## Acknowledgments

Z.J. was supported by the National Institutes of Health under Award Number R35GM154865.

## Author contributions

Z.J. conceived the study. H.W. and Z.J. conducted the analysis and wrote the manuscript.

## Competing interests

All authors declare no competing interests.

## Notes

### Competing Interest Statement

The authors have declared no competing interest.

### Summary of Updates

We have substantially revisited and expanded the analysis compared to the previous version.

